# Enumerating the Economic Cost of Antimicrobial Resistance Per Antibiotic Consumed to Inform the Evaluation of Interventions Affecting their Use

**DOI:** 10.1101/206656

**Authors:** Poojan Shrestha, Ben S Cooper, Joanna Coast, Raymond Oppong, Nga T. T. Do, Tuangrat Podha, Olivier Celhay, Philippe J. Guerin, Heiman Wertheim, Yoel Lubell

## Abstract

**Background:** – Antimicrobial resistance (AMR) poses a colossal threat to global health and incurs high economic costs to society. Economic evaluations of antimicrobials and interventions such as diagnostics and vaccines that affect their consumption rarely include the costs of AMR, resulting in sub-optimal policy recommendations. We estimate the economic cost of AMR per antibiotic consumed, stratified by drug class and national income level.

**Methods:** – The model is comprised of three components: correlation coefficients between human antibiotic consumption and subsequent resistance; the economic costs of AMR for five key pathogens; and consumption data for antibiotic classes driving resistance in these organisms. These were used to calculate the economic cost of AMR per antibiotic consumed for different drug classes, using data from Thailand and the United States (US) to represent low/middle and high-income countries.

**Results:** – The correlation coefficients between consumption of antibiotics that drive resistance in *S. aureus, E. coli, K. pneumoniae, A. baumanii*, and *P. aeruginosa* and resistance rates were 0.37, 0.27, 0.35, 0.45, and 0.52, respectively. The total economic cost of AMR due to resistance in these five pathogens was $0.5 billion and $2.8 billion in Thailand and the US, respectively. The cost of AMR associated with the consumption of one standard unit (SU) of antibiotics ranged from $0.1 for macrolides to $0.7 for quinolones, cephalosporins and broad-spectrum penicillins in the Thai context. In the US context, the cost of AMR per SU of antibiotic consumed ranged from $0.1 for carbapenems to $0.6 for quinolones, cephalosporins and broad spectrum penicillins.

**Conclusion:** – The economic costs of AMR per antibiotic consumed were considerable, often exceeding their purchase cost. Differences between Thailand and the US were apparent, corresponding with variation in the overall burden of AMR and relative prevalence of different pathogens. Notwithstanding their limitations, use of these estimates in economic evaluations can make better-informed policy recommendations regarding interventions that affect antimicrobial consumption and those aimed specifically at reducing the burden of AMR.

## INTRODUCTION

Human antimicrobial consumption, whether or not clinically warranted, is associated with propagation of antimicrobial resistance (AMR) [1, 2]. This and other key drivers of AMR are listed in Figure 1, notably widespread antibiotic use prophylactically and as growth promoters in agriculture [3].

**Figure 1.**
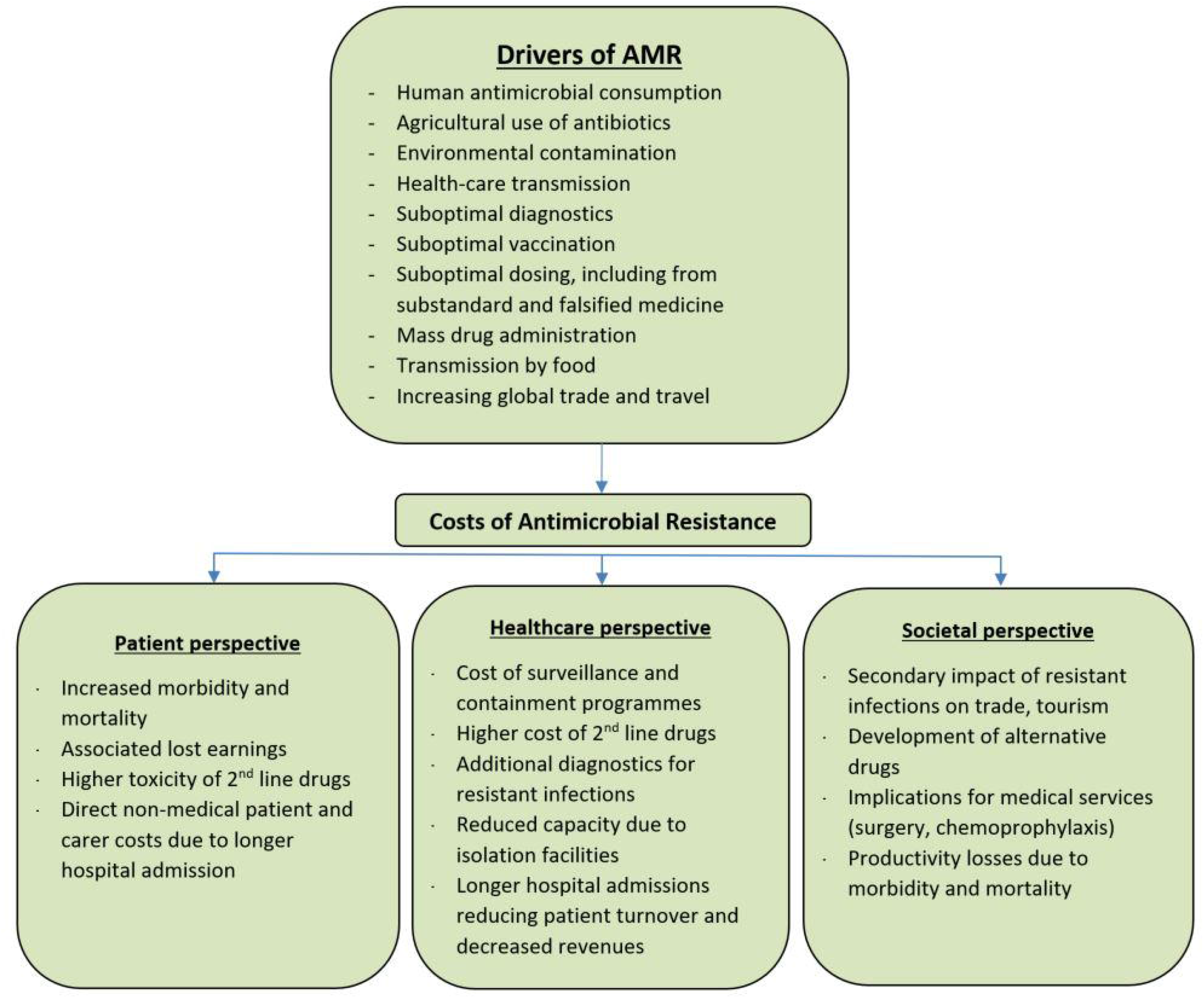
- Drivers and costs associated with antimicrobial resistance. Adapted: Holmes et al. [2] and McGowan [10]

Treatment of resistant infections is associated with higher costs for second line drugs, additional investigations, and longer hospitalisation [4]. Other indirect costs associated with AMR include productivity losses due to excess morbidity and premature mortality. These costs can be conceptualised as a negative externality to antimicrobial consumption accrued by all members of society, which are not reflected in the market price of antimicrobials [5, 6].

In addition to curative use in infectious diseases, antimicrobials are widely used presumptively, in mass treatment programmes (anti-helminths, antimalarials), and as prophylactics in surgical procedures and alongside immunocompromising treatments [2, 7]. Many other healthcare interventions such as vaccinations, diagnostics, and treatments for infectious diseases affect antimicrobial consumption, and consequently increase or decrease the risks of AMR. Economic evaluations of such interventions, however, have failed to internalise the potential costs of AMR into the analyses, leaving policymakers to intuitively consider these alongside more tangible costs and benefits in the evaluation [4, 8]. This can result in uninformed decision making, as the cost of AMR is likely to be under- or over-estimated by policymakers, if it is considered at all [4, 8, 9].

In 1996 Coast et al. argued that the omission of the cost of AMR in economic evaluation is partly explained by the challenges to its quantification [4], with extensive uncertainties surrounding resistance mechanisms, paucity and poor quality of relevant data, and other methodological challenges [5, 10]. The (mis)perception that the impact of AMR will only be felt in future years might also deter analysts from including them in the evaluation, assuming policymakers operate with a myopic view of health gains and costs. As confirmed in a recent review, very few attempts have since been made to quantify the externality of AMR [11].

Policymakers and key stakeholders, however, appear increasingly concerned with AMR, with unprecedented funding being allocated to interventions to mitigate its impact. In late 2016 the UN General Assembly held a special meeting on the topic, passing a unanimous resolution from Member States committing to adopt such measures [12]. Without enumerating the cost of AMR per antimicrobial consumed, it will be difficult to determine the allocative efficiency of these investments, and particularly so in low/middle income countries (LMICs) with more tangible causes of ill-health to invest in.

Therefore, despite the challenges, there is a clear need for costing the negative externality of AMR that can be affixed to the consumption of antimicrobials. The rare occasions where this has been done indicate the importance of such efforts. In a German hospital setting, for example, the use of a single defined daily dose of a second or third generation cephalosporin was associated with €5 and €15 respectively in costs of AMR [6]. The current analysis produced a menu of economic costs of AMR per antibiotic consumed for a variety drug classes, stratified into LMICs and high-income country settings. The output can be applied in future economic evaluations of interventions that involve or affect antibiotic consumption.

## Methods

### Economic costs of resistance

The economic cost of AMR is narrowly defined as the incremental cost of treating patients with resistant infections as compared with sensitive ones, and the indirect productivity losses due to excess mortality attributable to resistant infections. We therefore make a fundamental conservative assumption that resistant infections replace, rather than add to the burden of sensitive infections, even though there are strong indications that for Methicillin resistant *Staphylococcus aureus* (MRSA), for instance, the burden is additive to that of Methicillin sensitive *Staphylococcus aureus* (MSSA) [13]. We estimate these direct and indirect costs for the following key pathogens:

1. *Staphylococcus aureus* (*S. aureus*) resistant to Oxacillin
2. *Escherichia coli* (*E. coli*) resistant to 3^rd^ generation cephalosporin
3. *Klebsiella pneumoniae* (*K. pneumonia*) resistant to 3^rd^ generation cephalosporin
4. *Acinetobacter baumanii (A. baumanii)* resistant to carbapenems
5. *Pseudomonas aeruginosa (P. aeruginosa)* resistant to carbapenems

We focus our analysis on Thailand and the United States as representatives of low/middle and high-income country settings, respectively.

### Total economic loss

This is captured through the addition of the direct and indirect economic effects of AMR. The direct economic cost refers to the direct medical cost attributable to the treatment of a resistant infection as compared with the costs of treating a susceptible strain of the pathogen, and the indirect cost refers to the cost to society due to productivity losses attributable to premature excess deaths due to resistance.

#### Direct cost to the provider

We use the product of the number of resistant infections due to each of the above organisms, and the direct incremental medical cost attributable to resistance in the respective infections (Table 1). The number of infections and deaths per infection for the US was obtained from the Center for Disease Control and Prevention (CDC) [14]. The unit cost per infection was obtained from a study reporting the incremental cost of resistant bacterial infections based on the Medical Expenditure Panel Survey, with data available for 14 million bacterial infections of which 1.2 million were estimated to be antibiotic resistant [15]. These costs were inflation adjusted to 2016 US$ using the US consumer price index [16].

**Table 1.** – Incidence and mortality of resistant infections per 100,000, and the excess direct cost per resistant infection

|  | Mortality per 100,000 |  | Infections per 100,000 |  | Direct medical costs per infection |  |
| --- | --- | --- | --- | --- | --- | --- |
| | Thailand [18] | US [14] | Thailand | US | Thailand (US\$) [17] | US (US\$) [15] |
| *S.aureus* | 4.1 | 3.5 | 29.5 | 25.2 | 1,551 | 1,415 |
| *E. coli* | 0.9 | 0.2 | 13.3 | 3.3 | 956 | 1,415 |
| *K. pneumoniae* | 0.4 | 0.5 | 6.5 | 7.8 | 956 | 1,415 |
| *A. baumannii* | 22.4 | 0.2 | 326.9 | 2.3 | 1,749 | 1,415 |
| *P.aeruginosa* | 0.4 | 0.1 | 6.1 | 2.1 | 1,601 | 1,415 |
| **TOTAL** | **28.2** | **4.6** | **382.3** | **40.7** |  |  |

Estimates for the number of resistant infections and deaths in Thailand were available from two studies deriving their estimates from hospital records. The first report, published in 2012, estimated the number of AMR deaths at 38,000 [17], but we opted for the more conservative estimates in a 2016 study reporting approximately 19,000 AMR attributable deaths annually [18]. We obtained the unit cost per infection from the first of these studies, which included only the costs for antibiotics. We used an estimated excess length of stay (LoS) of 5 days for all gram negative bacteria based on the excess LoS for resistant *E. coli* infections [19] while for MRSA we assumed no excess LoS as compared with MSSA [20]. We then applied a cost of $38 per bed-day in a secondary hospital in Thailand to any excess LoS [21. 22], Costs were adjusted to 2016 US$ by converting to US$ at the year they were reported, and inflation adjusted using the World Bank Gross Domestic Product (GDP) deflator for Thailand.

#### Indirect cost

Mortality figures were converted into productivity losses taking the human capital approach, by multiplying them by an assumed ten productive life years lost per death, based on a study of survival post intensive care unit (ICU) admission in Thailand, which reported similar results for high income settings [22], with a sensitivity analysis of 5-20 productive years lost per death. The number of years lost was then multiplied by GDP per capita to generate the productivity losses per death. A 3% discount rate along with a 1% annual productive growth rate was applied to these values.

### Resistance modulating factor (RMf)

As illustrated in Figure 1, human antimicrobial consumption is one of a host of factors driving AMR, and different drug classes are implicated in propagating resistance in different pathogens. The Resistance Modulating factor (RMf) approximates the proportional contribution of human antimicrobial consumption towards the total cost of AMR. Correlation coefficients were calculated to study the strength of the relationship between consumption of antibiotic classes assumed to be implicated in driving resistance in each pathogen, and the rates of resistance observed to their first line treatments. It was assumed that drug classes that were implicated in driving resistance in each pathogen (Table 2) did so equally [23, 24]. Data points for consumption (from 2008 to 2014) and resistance (from 2008 to 2015) were obtained from 44 countries and included total consumption in both hospital and community settings [25].

**Table 2.**
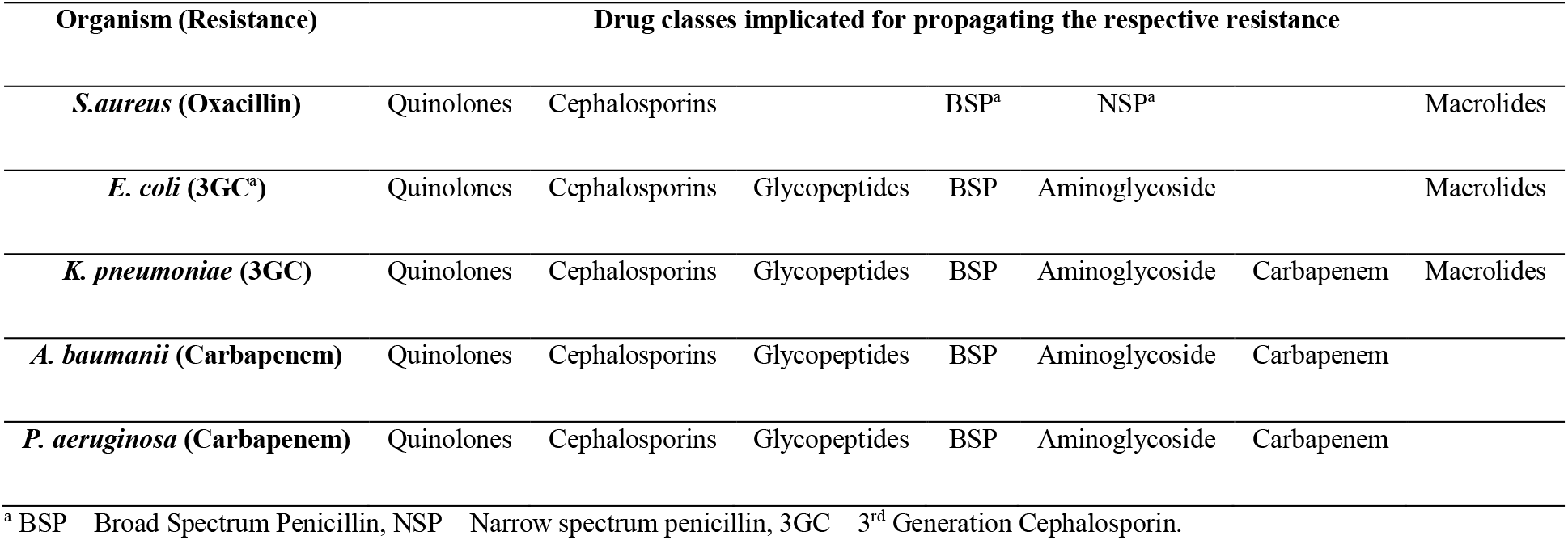
- Drug classes implicated in increasing the risk of resistance in each organism

| Organism (Resistance) | Drug classes implicated for propagating the respective resistance |  |  |  |  |  |  |
| --- | --- | --- | --- | --- | --- | --- | --- |
| *S.aureus* (Oxacillin) | Quinolones | Cephalosporins |  | BSP<sup>a</sup> | NSP<sup>a</sup> |  | Macrolides |
| *E. coli* (3GC<sup>a</sup>) | Quinolones | Cephalosporins | Glycopeptides | BSP | Aminoglycoside |  | Macrolides |
| *K. pneumoniae* (3GC) | Quinolones | Cephalosporins | Glycopeptides | BSP | Aminoglycoside | Carbapenem | Macrolides |
| *A. baumannii* (Carbapenem) | Quinolones | Cephalosporins | Glycopeptides | BSP | Aminoglycoside | Carbapenem |  |
| *P. aeruginosa* (Carbapenem) | Quinolones | Cephalosporins | Glycopeptides | BSP | Aminoglycoside | Carbapenem |  |

<sup>a</sup> BSP – Broad Spectrum Penicillin, NSP – Narrow spectrum penicillin, 3GC – 3<sup>rd</sup> Generation Cephalosporin.

The ecological association between the consumption of antibiotics implicated in driving resistance and the level of resistance was measured using Pearson’s correlation coefficient, *ρ_p_* for each pathogen *p*, considering the correlation between average resistance rates from 2008 to 2015 and the average of antibiotic consumption between 2008 and 2014. This is given by 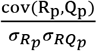 (Equation 1) where *R_p_* is the log transformed average annual measure of resistance for pathogen *p* (defined as the proportion of non-susceptible isolates), and *Q_p_* is the log-transformed mean consumption of implicated antibiotics. The denominators represent corresponding standard deviations. The lower and upper bounds of the 95% coefficient confidence intervals (CI) were used in the sensitivity analysis.

### Model for the economic cost of AMR per antibiotic consumed

Putting together the costs of AMR, the RMf, and the consumption of antibiotics that drive resistance in each pathogen, we established the cost of AMR attributable to the use of a Standard Unit (SU) and a full course of eight antibiotic drug classes. One SU is a measure of volume based on the smallest identifiable dose given to a patient, dependent on the pharmaceutical form (a pill, capsule, tablet or ampoule) [26]. The cost of AMR per SU is thus calculated as 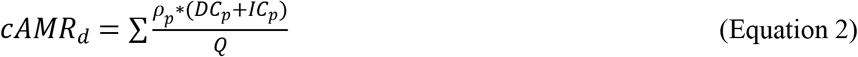 (Equation 2) where *cAMR* is the cost of AMR per standard unit of antibiotic *d* consumed, *DC* the direct cost of treatment and *IC* the indirect costs for each pathogen *p*, and *Q* is the annual consumption of antibiotics assumed to be implicated in driving resistance in the pathogen *p*. For each drug *d* the costs on the right of the equation are summed up for all pathogens in which it is implicated in driving resistance, as shown in Equation 2.

The resulting economic costs per SU of antibiotic consumed in each pathogen were then aggregated to calculate the cumulative economic cost per antibiotic consumed for each drug class in each country, including only the infections in which the particular drug class was assumed to propagate resistance. For example, as quinolones are assumed to drive resistance in all 5 pathogens the cost of resistance per SU of quinolones would be the sum of the cost of resistance shown in Equation 2 for all 5 pathogens.

Model outputs are also presented in terms of the cost of AMR per full course of treatment. While in reality there will be much variation in the number of SUs per course depending on the indication, patient age and other factors, we use a pre-specified number of SU per adult full course of antibiotics according to the British National formulary (BNF) [27]. The number of SU per full course ranged from 3 SU for a full course of macrolide antibiotics to 28 SU per full course of quinolones. The number of SUs per course for all classes is presented in Supplementary Table 1, Additional file 1.

#### Sensitivity analysis

The lower and the upper bound costs of AMR are calculated using the confidence intervals of the RMf and a range of 5-20 productive life years assigned to each excess death to calculate the indirect cost.

Data entry, verification, and analysis were done in Microsoft Excel 2016. Calculation of the correlation coefficients was done in R version 3.2.2 (R Foundation for Statistical Computing, Vienna, Austria). <u>A web interface for the model</u> where readers can vary parameter estimates and test model assumptions was developed using R-Shiny (RStudio, Boston, US) [28].

## Results

### The Resistance Modulating factor

As shown in Table 3, a positive relationship was confirmed between consumption of antibiotics assumed to be implicated in resistance, and the average resistance rates in all pathogens, with correlation coefficients ranging from 0.27 in *E. coli* (p=0.07) to 0.52 in *P. aerginosa* (p=0.0006).

**Table 3.** - Pearson’s correlation coefficient showing ecological associations between average consumption (2008-14) and corresponding resistance (2008-15)

| Organism / resistance | Correlation coefficient (95% CI, p-values) |
| --- | --- |
| *S. aureus* resistant to oxacillin | 0.37 (0.08 - 0.61, p = 0.016) |
| *E. coli* resistant to 3rd generation cephalosporin | 0.27 (-0.03 - 0.53, p = 0.07) |
| *K. pneumoniae* resistant to 3rd generation cephalosporin | 0.35 (0.06 - 0.59, p = 0.019) |
| *A. baumannii* resistant to carbapenem | 0.45 (0.15 - 0.68, p = 0.005) |
| *P. aeruginosa* resistant to carbapenem | 0.52 (0.25 - 0.72, p = 0.0006) |

### Direct and indirect costs of AMR

The total economic cost of AMR due to drug resistance in the five pathogens was $0.5 billion and $2.8 billion in Thailand and the United States, respectively. This is disaggregated into direct and the indirect costs for each of the organisms in the two countries in Tables 4 and 5, respectively. As an illustration, the direct and indirect annual cost of AMR in Thailand due to MRSA was estimated at $29 million and $151 million, respectively. After adjusting for the relative contribution of human consumption using the RMf, the direct and indirect economic loss was estimated to be $11 million and $56 million.

### Economic cost of AMR per antibiotic consumed

With the total economic cost of AMR for each pathogen multiplied by its RMf in the numerator, and the consumption data for the relevant drug classes in the denominator, the economic cost of AMR of one SU of antibiotic for each pathogen was calculated (Table 6). Thus any antibiotic implicated in driving resistance in *S. aureus* (Table 2) would have an economic cost of AMR of $0.07 per SU in the Thai setting, and if a full course of the same drug consisted of 10 units this would imply a cost of $0.69 per full course.

**Table 5.** – Productivity losses due to excess deaths attributable to resistant infection (Indirect Cost)

|  | Thailand |  |  |  |  | United States |  |  |  |  |
| --- | --- | --- | --- | --- | --- | --- | --- | --- | --- | --- |
|  | *S. aureus* | *E. coli* | *K. pneumoniae* | *A. baumannii* | *P. aeruginosa* | *S. aureus* | *E. coli* | *K. pneumoniae* | *A. baumannii* | *P. aeruginosa* |
| Excess deaths | 2,799 | 597 | 288 | 15,168 | 270 | 11,285 | 690 | 1,620 | 500 | 440 |
| GDP/capita (US\$)<sup>a</sup> | | | 5907 | | | | | 57466 | | |
| Indirect Cost (million US\$) | 151 | 32 | 16 | 815 | 15 | 5,901 | 361 | 847 | 262 | 230 |
| RMf | 0.37 | 0.27 | 0.35 | 0.45 | 0.52 | 0.37 | 0.27 | 0.35 | 0.45 | 0.52 |
| Indirect cost due to human consumption (million US\$) | 56 | 9 | 5 | 367 | 8 | 2,184 | 97 | 297 | 118 | 120 |

<sup>a</sup> Data from World Bank

As most antibiotics are assumed to drive resistance in more than one infection, the costs need to be aggregated for all relevant pathogens to obtain the cumulative cost of AMR attributable to the consumption of one SU of that antibiotic. For a broad spectrum penicillin that is assumed to drive resistance in all pathogens, the estimated cost of AMR would be $6.95 per course of 10 SU in Thailand. The costs in Table 6 were therefore aggregated for each drug class where it was assumed to drive resistance in each of the organisms. Table 7 presents the cumulative economic cost per SU and per full course by drug class.

**Table 6.**
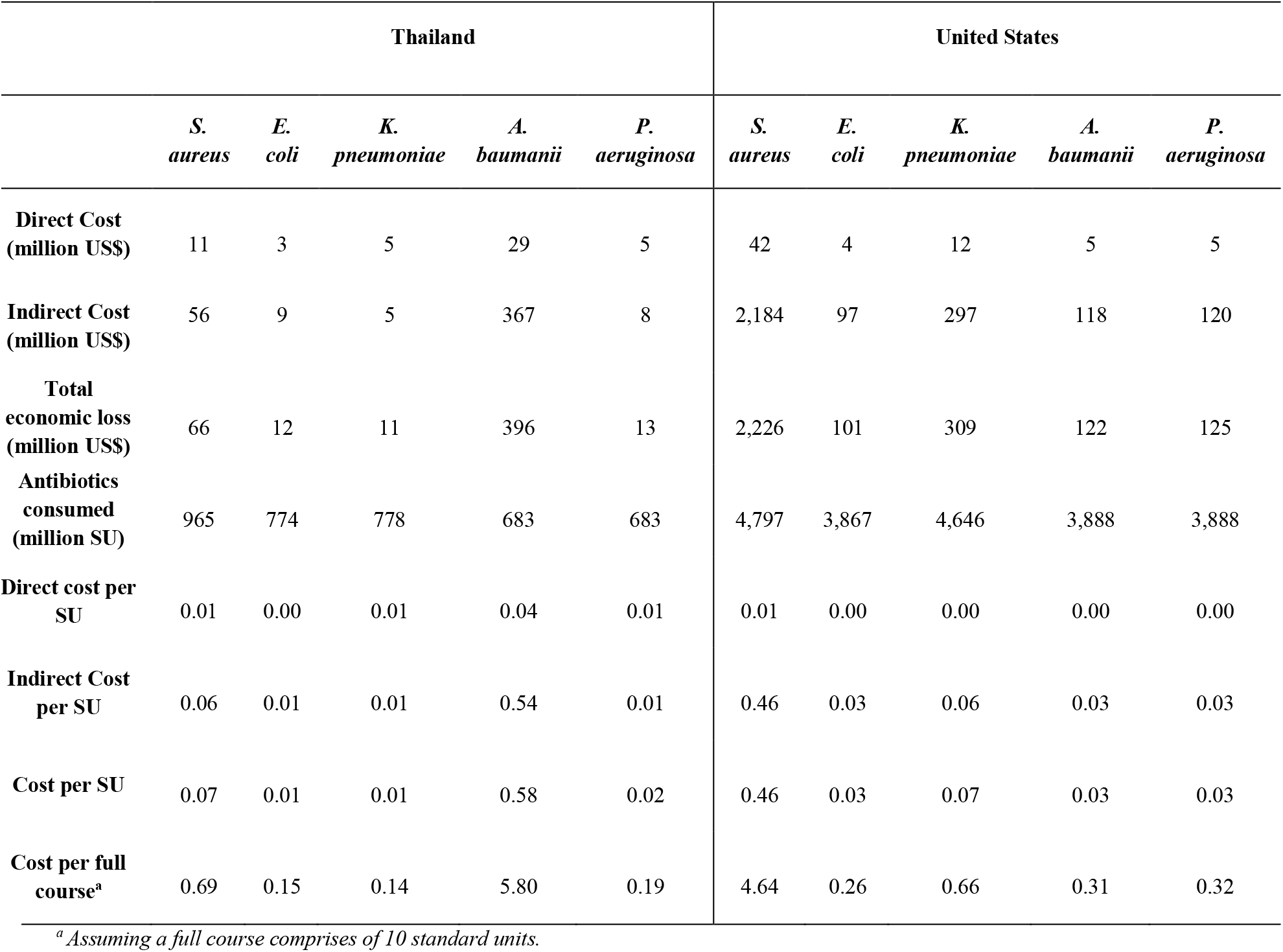
- Cost per Standard Unit (SU) and full course antibiotic consumed per resistant organism

**Table 7.**
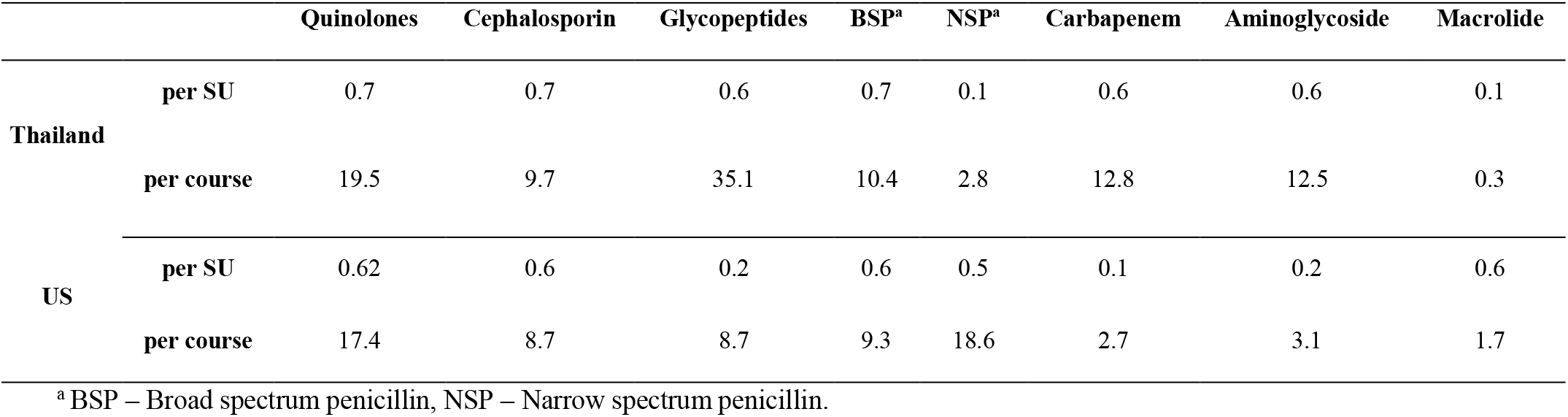
- Cumulative cost per SU and per antibiotic course by drug class (US$)

### Sensitivity analysis

The lower and the upper bound costs of AMR were calculated using the confidence intervals of the RMf (Table 3) and a range of 5-20 productive life years assigned to each excess death for the indirect cost of AMR. Table 8 shows the resulting range of economic costs for a SU and a full course of antibiotic consumed in Thailand and US. Hence, in Thailand, the best case scenario would see a cost of AMR of $2.93 per course of co-amoxiclav and the worst would be $32.16.

**Table 8.** - Range of economic costs per full course of antibiotics using outputs from the sensitivity analysis (US$).

| Antibiotic (Drug class) | Thailand |  | United States |  |
| --- | --- | --- | --- | --- |
|  | Cost per SU | Cost per full course | Cost per SU | Cost per course |
| Levofloxacin (Quinolone) | 0.2 – 2.1 | 5.5 – 60.0 | 0.1 – 1.8 | 2.1 – 51.2 |
| Ceftriaxone (Cephalosporin) | 0.2 – 2.1 | 2.7 – 30.0 | 0.1 – 1.8 | 1.0 – 25.6 |
| Vancomycin (Glycopeptide) | 0.2 – 2.0 | 10.4 – 109.4 | 0.0 – 0.5 | 1.1 – 25.5 |
| Co-amoxiclav (BSP) | 0.2 – 2.1 | 2.9 – 32.2 | 0.1 – 1.8 | 1.1 – 27.4 |
| Phenoxymethylpenicillin (NSP) | 0.0 – 0.2 | 0.4 – 7.6 | 0.1 – 1.4 | 2.2 – 54.9 |
| Meropenem (Carbapenem) | 0.2 – 1.9 | 3.9 – 40.1 | 0.0 – 0.4 | 0.4 – 7.6 |
| Amikacin (Aminoglycoside) | 0.2 – 2.0 | 3.7 – 39.1 | 0.0 – 0.5 | 0.4 – 9.1 |
| Azithromycin (Macrolide) | 0.0 – 0.3 | 0.0 – 0.8 | 0.1 – 1.7 | 0.2 – 5.0 |

## Discussion

Evidence-based policy draws on economic evaluation to allocate resources most efficiently [29], but this is entirely dependent on the inclusion of all pertinent costs and benefits associated with interventions under consideration. This is, to our knowledge, a first attempt at estimating the costs of AMR per antibiotic consumed by drug class and across national income brackets. We chose simple and transparent methods and restricted our assessment to the current burden of AMR, rather than more uncertain future projections, and to tangible factors including only direct medical costs and productivity losses due to AMR attributable deaths. Even within this restrictive framework there is much uncertainty surrounding interactions between antibiotic consumption, development of resistance, and its economic implications, but our underlying assumptions and parameter estimates were conservative.

The cost per SU of antibiotic differed between the US and Thailand for several reasons. First, the burden of AMR is considerably higher in Thailand, with a total of 28 AMR associated deaths per 100,000 as compared with 4.6 per 100,000 in the US (Table 1). Furthermore, the two countries had different epidemiological profiles, such as a higher burden of *Acinetobacter* associated mortality in Thailand as compared with the dominance of MRSA in the US. There were also notable differences in the cost data between the two countries; as the unit costs per infection for Thailand were only available from hospital settings, they tended to be higher than those in the US, which included both hospital and community settings. Other factors contributing to this difference are the higher GDP per capita and lower per capita consumption of antibiotics in the US.

The costs of AMR for drug classes also varied widely, driven primarily by the degree to which they were assumed to propagate resistance in the selected infections; NSPs were assumed to drive resistance only in *S. aureus*, while cephalosporins were implicated in resistance in all pathogens. The costs per full course of antibiotics were mostly determined by the number of SU per course, which for glycopeptides is high - a full course of vancomycin being 56 SU (four daily over 14 days) as compared with three daily units for a course of azithromycin (Supplementary Table 1, Additional file 1).

Very few attempts have been made to quantify the cost of AMR per antibiotic consumed and internalise them in evaluations of interventions that involve or affect the use of antimicrobials. A recent study by Oppong et al. was one of the first attempts to do so in an evaluation focusing on antibiotic treatment of respiratory infections, demonstrating the decisive impact this had on outcomes [30]. Their estimate for the cost of AMR, however, assumed that resistance is driven exclusively by human antimicrobial consumption and that consumption of all drug classes contribute to resistance in all pathogens equally. It also ignored the considerable differences in the burden of resistance across countries, as apparent in the much higher burden of AMR in Thailand compared with that in the US. An earlier study evaluating the cost-effectiveness of malaria rapid tests used a similarly crude estimate for the cost of antimalarial resistance, also showing the large impact this had in swaying results and conclusions [31]. Elbasha, building on previous work by Phelps [32] estimated the deadweight loss of resistance due to overtreatment and found a higher cost of AMR of $35 (2003) per course of amoxicillin in the US context [33].

Several studies have explored the correlation between antimicrobial consumption and resistance [34–36]. The correlation coefficients in the current study are smaller than prior estimates. For example, the coefficient for resistance in *E. coli* in this analysis was 0.27 (Table 4) in comparison to 0.74 from Goossens et al. [34]. This could be explained by the latter using 14 European countries in contrast to 44 countries from different regions in our study, and more abundant data for European countries that enabled correlating between the consumption and resistance of specific drugs, rather than drug classes as done here. The smaller coefficients imply a conservative assessment of the cost of AMR attributable to human antibiotic consumption.

**Table 4.** – Direct cost to the providers due to human antibiotic consumption in each resistant infection.

|  | Thailand |  |  |  |  | United States |  |  |  |  |
| --- | --- | --- | --- | --- | --- | --- | --- | --- | --- | --- |
|  | *S. aureus* | *E. coli* | *K. pneumoniae* | *A. baumannii* | *P. aeruginosa* | *S. aureus* | *E. coli* | *K. pneumoniae* | *A. baumannii* | *P. aeruginosa* |
| **Total infections** | 18,725 | 11,116 | 15,239 | 36,553 | 6,118 | 80,461 | 10,400 | 24,900 | 7,300 | 6,700 |
| **Cost per infection** | 1,551 | 956 | 956 | 1,749 | 1,601 | 1,415 | 1,415 | 1,415 | 1,415 | 1,415 |
| **Direct cost (million US\$)** | 29.0 | 10.6 | 14.6 | 63.9 | 9.8 | 113.8 | 14.7 | 35.2 | 10.3 | 9.5 |
| **RMf** | 0.37 | 0.27 | 0.35 | 0.45 | 0.52 | 0.37 | 0.27 | 0.35 | 0.45 | 0.52 |
| **Direct cost due to human consumption (million US\$)** | 10.7 | 2.9 | 5.1 | 28.8 | 5.1 | 42.1 | 4.0 | 12.3 | 4.6 | 4.9 |

Kaier et al. derived measures of association between antibiotic consumption and resistance from a time-series analysis using a multivariate regression model with different drug classes [37]. This would be a better approach for calculating the RMf, rather than the ecological associations used here. We were restricted, however, by having only 10 years of consumption data and even sparser and more heterogeneous resistance data.

There were many assumptions and limitations in the analysis (see Supplementary Table 2, Additional file 1). One key limitation was the inclusion of a limited number of organisms, while consumption of the same antibiotics could drive resistance in other organisms with additional costs. The Thai estimates also focused only on the burden of AMR within hospital settings, excluding the possible excess burden in primary care and the community. These and other listed limitations result in a conservative estimate of the economic costs of AMR in our model.

Taking the human capital approach to productivity losses implies much higher estimates than would have been derived using friction costs; given the context of this analysis, trying to capture the full societal costs of AMR, this was deemed appropriate. This is essentially equivalent to the widespread use of GDP/capita as a proxy for the ceiling ratio in cost-effectiveness analyses to classify interventions as cost-effective.

The direct medical costs assigned to resistant infections were derived very differently in each country; the US estimates were taken from a recent study providing a national estimate of the incremental healthcare cost of treating millions of patients with antibiotic sensitive and resistant infections [15]. The Thai estimates used rudimentary costing methods, largely relying on expert opinion to estimate the cost of antibiotics required to treat resistant infections.

The selection of drug classes implicated in propagation of resistance in the respective organisms were based on limited available evidence [24]. This might explain some apparent anomalies, like the relatively low costs for NSPs, which were assumed to drive resistance only in *S. aureus*. Another reason for this anomaly relates to the entire framework of the analysis, whereby the cost of AMR is approximated from its current (or recent) estimated burden, rather than projections of what will happen if resistance to last line drugs, such as carbapenem, were to spread, for which there are alarming early indications. Such an approach is arguably more relevant than focusing on the present burden of AMR, but it requires many more strong and contestable assumptions.

The data on consumption and resistance levels used to derive the RMf were limited to 10 years and a causal relationship was assumed. For many pathogens and types of infections, however, this is not realistic as increasing resistance could alter consumption patterns as patients and physicians adapt their behaviour in order to provide the best possible treatment in a changing environment of resistance and therefore counteract the assumed dose-response relationship.

These rudimentary estimates for the economic cost of AMR per antibiotic consumed could be improved upon in several ways in future work as better data become available. In addition to addressing the above limitations, the link between human antibiotic consumption and resistance can be disaggregated into hospital vs. community use. The model can be further extended to other organisms including parasites and viruses and their varying distribution in different health sectors and geographical locations (global/regional/country/hospital/community).

## Conclusions

The estimates of the economic costs of AMR per antibiotic consumed in this analysis were high. Incorporation of such estimates in economic evaluation of interventions that affect the use of antibiotics will better portray their true costs and benefits, and could act as a catalyst for more efficient deployment of interventions to mitigate the burden of AMR. We highlight the limitations of the analysis to emphasise the need for further development of the methods, and point to the notable differences in the costs of AMR per antibiotic consumed between the two countries and within the different drug classes to encourage their adaptation to other settings as relevant data become available.

## Declarations

### Ethics approval and consent to participate

Not applicable

### Consent for publication

Not applicable

### Availability of data and materials

The antibiotic consumption (2008 - 2014) and pathogen resistance (2008-2015) data used in this study was obtained from https://resistancemap.cddep.org/ and is openly accessible. [25]

### Authors’ contribution

YL and PS conceptualised and designed the study. PS, YL and BC analysed and interpreted the data. PS and YL drafted the manuscript. JC, RO, OC, NDTT, TP, PG and HW revised the manuscript for intellectual content. OC designed the web-interface. All authors read and approved the final manuscript.

## Acknowledgements

We thank Ms. Nistha Shrestha for her contribution in the data compiling process. We also thank Professor Lisa White, Dr Pan-Ngum Wirichada and other members of the Mathematical and Economic Modelling group at the Mahidol Oxford Tropical Medicine Research Unit for their helpful feedback for a presentation of this analysis.

## List of Abbreviations

AMR: - Antimicrobial resistance
BNF: - British National formulary
CDC: - Center for Disease Control and Prevention
CI: - confidence interval
GDP: – Gross Domestic Product
ICU: – Intensive Care Unit
LMICs: - low/middle income countries
LoS: – Length of Stay
MRSA: - Methicillin resistant *Staphylococcus aureus*
MSSA: - Methicillin sensitive *Staphylococcus aureus*
RMf: - Resistance modulating factor
SU: - Standard Unit
US: - United States

